# Inexpensive, scalable camera system for tracking rats in large spaces

**DOI:** 10.1101/285460

**Authors:** Rajat Saxena, Warsha Barde, Sachin S. Deshmukh

**Author notes:** **Correspondence** Sachin S. Deshmukh, Centre for Neuroscience, Indian Institute of Science, Bangalore. **Author Contributions:** SSD conceived the design. RS assembled the system and wrote the codes. WB and RS performed the experiments and collected the data. RS analyzed the data. RS and SSD wrote the manuscript.

## Abstract

Most studies of neural correlates of spatial navigation are restricted to small arenas (≤ 1 m^2^) because of the limits imposed by the recording cables. New wireless recording systems have a larger recording range. However, these neuronal recording systems lack the ability to track animals in large area, constraining the size of the arena. We developed and benchmarked an open-source, scalable multi-camera tracking system based on low-cost hardware. This camera system was used in combination with a wireless recording system for characterizing neural correlates of space in environments of sizes up to 16.5 m^2^. This system improved accuracy in estimating spatial firing characteristics, theta phase precession, and head direction tuning of neurons compared to a popular commercial system, due to its better temporal accuracy. This improved temporal accuracy is crucial for accurately aligning videos from multiple cameras in large spaces and characterizing spatially modulated cells in a large environment.

## Introduction

Spatial navigation is a widely employed behavior to study the neuronal circuits underlying cognition, learning, and memory. Since the discovery of place cells in the hippocampus four decades ago (O’Keefe & Dostrovsky, 1971; O’Keefe & Nadel, 1978), a considerable amount of work has been undertaken to understand the representation of space in the brain. Over the years, a variety of cell types such as grid cells (Hafting et al., 2005), head direction cells (Ranck, 1984; Taube et al., 1990), speed cells (Kropff et al., 2015), border cells (Savelli et al., 2008; Solstad, et al., 2008), object cells (Deshmukh & Knierim, 2011), and landmark vector cells (Deshmukh & Knierim, 2013) have been recorded from the hippocampal formation, advancing our understanding of its role in spatial navigation.

Because of the limits imposed by the cables extending from the animal to the recording system, these experiments studying spatial maps have largely been limited to small spaces (≤ 1 m^2^), with rare exceptions (for example, 1.5 m × 1.4 m in Fenton et al., 2008; 1.8 m × 1.4 m in Park et al., 2011; 2.2 m × 2.2 m in Stensola et al., 2012; 3.5 m diameter circular arena in Gothard et al., 1996; 18 m track in Kjelstrup et al., 2008; 48 m track in Rich et al., 2014). This experimental constraint leaves a significant lacuna in our understanding of the neural correlates of spatial navigation in environments of scale and complexity comparable to the natural habitat of the rat. Home ranges of Norway rats vary from tens of m^2^ in urban areas to hundreds of m^2^ in farms and fields (Lambert et al., 2008; Oyedele et al., 2015); rat burrows occupy an area of the order of 10 m^2^ (Calhoun, 1963).

The advent of wireless recording systems enables us to now record neural activity from the hippocampal formation while the rat forages in larger and more complex environments. Most commercial neuronal recording systems, however, do not have provisions for recording from more than two cameras which again constrains the size of the behavioral arena, leading to an increased need for a system capable of tracking animals in larger environments. Using wide angle lenses with the standard cameras is not a satisfactory solution, as the resolution decreases drastically. Wide angle lenses with higher resolution 4K cameras can alleviate the resolution issue, but occlusion by the experimenter and the environmental features, cost, and synchronization with neural recording system make this solution sub-optimal.

Here, we describe a novel tracking system comprising 8 overhead Raspberry Pi cameras (referred to as the “Picamera system” henceforth), capable of tracking an animal’s position in a large environment. To benchmark the Picamera system, we compared its performance with a commercial video tracking system sold as a part of a wireless electrophysiology system. We recorded different cell types from the hippocampal formation using both these video trackers coupled with the wireless electrophysiology system. We show that the higher temporal accuracy of the Picamera system improved our ability to estimate multiple spatial firing characteristics of spatially modulated cells in standard environments used in spatial navigation studies. We then went on to record from a 5.5 m × 3 m arena using the Picamera system coupled with the wireless electrophysiology system to demonstrate our ability to characterize neural correlates of spatial navigation in a large space.

## Materials and Methods

### Animals and Surgical Procedures

Four male Long–Evans rats aged 5-8 months, weighing 450-600 gm were housed individually on a 12:12 hr reversed light/dark cycle and habituated to daily handling over 2 weeks before surgery. All experiments were performed according to a protocol approved by the Institutional Animal Ethics Committee of the Indian Institute of Science.

Custom-built hyperdrives having 16 tetrodes + 2 references were implanted over the right hemisphere under surgical anesthesia (80 mg/kg ketamine + 10 mg/kg xylazine followed by 0.5-2% isoflurane for maintenance). The tetrodes targeted different parts of the hippocampal formation in different rats: area CA1 of the hippocampus (2 rats), medial entorhinal cortex (MEC) (1 rat), MEC and Lateral Entorhinal Cortex (LEC) (1 rat).

### Training and Experimental Protocol

The rats recovered for 5-7 days after the surgery until their weights stabilized. During subsequent training and recordings, the rats were food deprived and trained to forage for food on a 1 m × 1 m black square platform. The neural recordings were performed in multiple setups: a 5.5 m × 3 m room (henceforth referred to as ‘the large room’), half of this same room, 2.75 m × 3 m, a circular track (diameter = 1 m), a linear track (length = 1 m) and the square platform used for training. Neural signals were recorded wirelessly in all these setups. For small setups (circular track, linear track, and square platform), a single Picamera sub-unit and a commercial camera were used for position tracking while the rats foraged for food. For the large room recordings, 8 Picamera sub-units covering the entire room with substantial inter-camera overlap were used for position tracking while the rats foraged for food.

### Neural Data Recording Hardware

Recordings were performed using the Cheetah data acquisition system and Cube-64, a 64-channel wireless transmitter (Neuralynx, Bozeman, MT, USA). A control computer ran the data acquisition software and stored the acquired data (Figure 1a).

**Figure 1:**
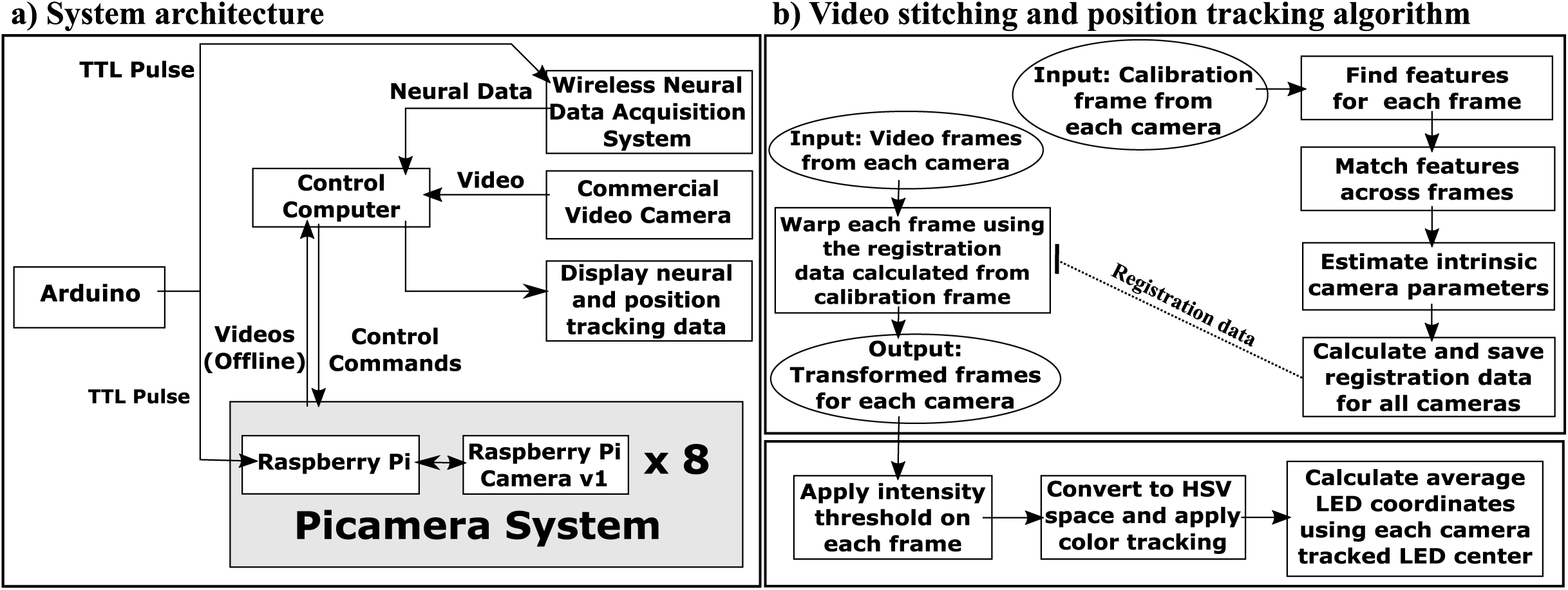
Recording setup. (a) Neural data acquired using the wireless data acquisition system and the video frames monitoring the animal are stored on the control computer. The Neuralynx Cheetah software processes, displays, and stores neural data as well as positions of tracking LEDs on the animal’s head. The Picamera system comprises 8 sub-units, each with a Raspberry Pi and a Raspberry Pi camera. Videos recorded from the Picamera are displayed on a monitor, and stored locally on each Raspberry Pi. The control computer is used to control all the Picamera sub-units and at the end of the session, receives the video and frame timestamp information acquired by the Picamera sub-units for longer term storage. An Arduino is programmed to send time sync signals (TTL pulses) to all the Picamera sub-units and the data acquisition system simultaneously (b) Top, calibration frames from all the cameras are loaded. Scale and rotation invariant features are extracted from each frame and matched across cameras. Using geometrically consistent feature matches, intrinsic camera parameters are estimated followed by calculation of registration data for each camera which is then saved. Each frame in the video from each camera is then warped and transformed to align it to a single coordinate reference frame using the saved registration data. Bottom, intensity thresholding followed by color tracking in HSV color space is applied on each camera frame to get the coordinates of red and green LEDs. The final position for each frame is estimated after averaging the positions from each camera.

### Video Cameras

The tracking system comprised 8 overhead Picamera sub-units (Figure 1a). Each sub-unit had a Raspberry Pi camera module v1 (containing an OmniVision OV5647 ColorCMOS QSXGA 5 MP sensor with f/2.9 aperture lens) connected to a Raspberry Pi 2 model B computer (1 GB RAM, 900 MHz quad-core processor and 32 GB class 10 SD card) (www.raspberrypi.org). We recorded video on each Picamera sub-unit at 640 × 480 resolution at 30 Hz with high temporal accuracy and no dropped frames over 4 hrs of recording. The Picamera system could acquire at higher frame rates with high temporal accuracy, but there were occasional frame drops (0.22% for 50 Hz and 0.31% for 60 Hz), and thus we chose to acquire at 30 Hz. Each camera has a field of view of 2.43 m × 1.82 m when mounted at a height of 2.8 m. The parallel architecture of the Picamera system, with each Raspberry Pi camera recording video on its own Raspberry Pi computer independently of others, makes it easily scalable without compromising temporal accuracy. The Picamera Python library (https://picamera.readthedocs.io/) was used to acquire video data along with timestamps from the cameras. Video recordings from the Picamera were displayed using a composite video output port available on each Raspberry pi.

The Picamera was compared to the standard camera kit supplied by Neuralynx Inc. for benchmarking. The camera kit (henceforth referred as ‘commercial camera’) comprised a CV-S3200 analog color camera with a 6 mm lens (JAI Inc., San Jose, CA, USA) and a HVR 1850 PCIe frame grabber card (Hauppauge Inc., Hauppauge, NY, USA) which is compatible with the Cheetah Software. The camera covered an area of 3 m × 2.3 m when mounted at a height of 2.8 m. The commercial camera recorded videos at 25 Hz frame rate due to the PAL encoding standard followed for India’s 50 Hz frequency of AC power (the NTSC encoding standard, which records videos at 30 Hz and is routinely used in countries with 60 Hz AC power supply, distorted the video in our setup). Since the Picamera system runs on 5 V DC power, it is not affected by the line frequency while acquiring at 30 Hz frame rate. To account for the potential confound due to different frame rates, we compared data recorded in one rat by acquiring video at 25 Hz on both the Picamera and the commercial camera.

### Time Synchronization across multiple Picamera sub-units and the data acquisition system

Each Raspberry Pi was connected via a DGS-1024D ethernet hub (D-Link, Taipei, Taiwan) to the control computer and remotely operated over the network using PuTTY (http://www.putty.org/) for secure shell (ssh).

In order to synchronize position tracking data (see below) acquired in these videos with neural recordings, it is critical to know the time at which each frame was recorded. Default video capture protocols do not record timestamps for each frame. The Picamera Python library offers an option to save operating system clock time, but this option is prone to jitter introduced by other processes competing for CPU resources. To reduce the jitter in frame timestamps, we modified the Picamera library. Since the Raspberry Pi’s Graphics Processing unit (GPU) runs its own real-time operating system, it allows saving frame timestamp with extremely low jitter. A Custom video output encoder was written to save the Raspberry Pi’s system time clock (STC) value acquired by the GPU each time the Raspberry Pi camera sent a “start of frame” interrupt signal to the GPU. This “presentation timestamp” accurately times each frame of the video.

TTL (Transistor-Transistor Logic) pulses generated using Arduino Uno REV3 (https://store.arduino.cc/usa/arduino-uno-rev3) were sent to all 8 Picamera sub-units as well as the data acquisition system for synchronizing time across video and neural data (Figure 1a). Timestamps for each TTL input on/off transition were logged on all the Raspberry Pi computers and the data acquisition system. Difference between timestamps recorded on each Raspberry Pi computer and the data acquisition system corresponding to the first TTL on transition gives us the instantaneous temporal offset between these devices. This offset was then subtracted from all subsequent frame timestamps of each video to convert their timestamps to the data acquisition system’s temporal reference frame. Subsequent TTL on/off transitions showed that there was virtually no temporal drift between the Raspberry Pi computers and the data acquisition system for up to 4 hrs of recording (in case of temporal drift/jump between clocks on different systems, every TTL on/off transition can be used to correct the errors).

Three files were generated on each Picamera sub-unit: a video file in .h264 format and two csv files: one containing the timestamps for all the frames and other one holding the timestamps corresponding to each TTL input on/off transition.

### Video Stitching

Videos from each camera were processed offline. A representation of the 5.5 m x 3 m room was created by aligning simultaneously captured video frames from all 8 Picamera sub-units, each of which covered only a part of the large room and had a substantial overlap with at least two cameras. Since we had the exact timestamp for each frame, we could temporally align each frame accurately across cameras. Video stitching involved two steps: 1. Calculation of registration data for each camera, and 2. Generation of aligned images for each camera (Figure 1b).

### Calculating Registration Data

We used Brown & Lowe’s (2007) algorithm to stitch frames from multiple cameras into a single representation. Briefly, one calibration frame (Supplementary Figure 1) was acquired from each camera once at the start of each experiment by laying down cloth/paper with uniquely patterned background to increase the number of identifiable features in the large room. We extracted feature vectors which are invariant to image scale, rotation and are also robust to changes in illumination, noise, and changes in viewpoints for calibration frames from all cameras. These features were matched across all cameras to find the nearest neighbor pixels between two or more cameras. The Random Sample Consensus (RANSAC) algorithm (Fischler & Bolles, 1981) was then applied to find geometrically consistent feature matches and generate registration data for each camera. The registration data for each camera included information about focal lengths (f_x_, f_y_), lens displacement along X and Y axis (c_x_, c_y_), radial and tangential distortion coefficients, rotation, and translation matrices. It is important to note that the registration data needs to be recalculated if the cameras are displaced/rotated from their original location.

### Generating the aligned Image for each camera

Individual video frames from all the cameras were then transformed using the registration data (calculated once at the beginning of experiment using one calibration frame from each camera) to get them all in the same coordinate system. We modified the OpenCV library Stitcher class (https://opencv.org/) to work with video files. The original OpenCV module code generates a single stitched image from the set of simultaneously recorded input images. In a single merged output frame, the intensity of LEDs used for tracking the rat would get averaged down, possibly below threshold used in position tracking (see below) when an experimenter or other objects occlude field of view of one camera but not another. We modified the code to generate transformed frames from the video of each camera rather than generating a single merged output frame. The individual camera frame transformation enabled us to tackle the problem of occlusion by experimenter as well as other objects by allowing us to perform thresholding operations on un-averaged image followed by averaging the positions estimated by individual cameras (see below). When the LEDs were visible on multiple cameras, individual camera frame transformation would perform as well as the single merged output frame.

### Position Tracking

Red and green LEDs mounted on the animal’s head allowed tracking the animal’s position and head direction. For each camera frame, all the pixels crossing a predefined intensity threshold were converted to the Hue, Saturation, Value (HSV) color space to identify red and green pixels. The appropriate pixel clusters were identified using a custom HSV color range for red (H = 0:10,160:180 S = 100:255 V = 50:255; OpenCV uses H range of 0-180 instead of the standard 0 to 360) and green (H = 50:70 S = 50:255 V = 100:255) colors and the coordinates of the center of the red and green pixel clusters were noted as the animal’s position for that frame. The animal’s final position corresponding to a frame was calculated by averaging the position estimate across each camera frame for both, the red and green LED lights (Figure 1b). Head direction of the animal was calculated using the positions of red (anterior) and green (posterior) LEDs.

### Data Analysis

#### Cluster Cutting

Manual spike sorting with a custom software (WinClust, J.J. Knierim, Johns Hopkins University) was employed to segregate spikes of isolated single units. Each unit was assigned an isolation quality score on a scale of 1 (very well-isolated) to 5 (poorly isolated) based on the separation of its cluster from other units and the background together with its inter-spike interval histogram. A cluster with an isolation quality of 1-3 and firing at least 50 spikes in a session was used for the subsequent analyses. Putative interneurons (mean firing rates > 10 Hz; Frank et al., 2001) were excluded.

#### Spatial Firing Rate Maps and Place Fields

The position data along with the spike counts were segmented into 4 cm × 4 cm spatial bins for the large room and 2 cm × 2 cm spatial bins for other setups. Times during which the rat moved < 2 cm/s and the spatial bins where the rat spent < 0.4 s were excluded from the analysis. The firing rate map of each cell was calculated by dividing the number of spikes fired in each bin by the time spent there. Rate maps smoothed using the adaptive binning algorithm (Skaggs et al., 1996) were used to calculate spatial information score (see below). Gaussian (sigma = 1.25 bins) smoothed rate maps were used to calculate peak firing rate, place field size and for illustrations. Only place fields with peak firing rate greater than the 25% of the peak firing rate for that cell’s rate map were included for the place field analysis. The size of an individual place field was determined as number of contiguous pixels (minimum 7) with firing rate greater than 15% of the peak firing rate of that field.

#### Head Direction

Head direction tuning curves were calculated after dividing the total number of spikes fired for each head direction bin (5° bin width) by the amount of time the rat spent facing in that angular bin (Taube et al., 1990) and smoothed with a Gaussian with sigma = 1.25 bins.

#### Spatial Information

A spatial information score (Skaggs et al., 1996) was used to quantify the spatial tuning of single units. The score calculates the information (in bits) about the rat’s location conveyed by a single spike. We employed a shuffling procedure to estimate the probability of obtaining the observed spatial information by chance. The spike train was shifted cyclically with respect to position data one thousand times by adding a uniformly generated random number lying between 30 seconds and the duration of the recording session - 30 seconds. The fraction of time-shifted information scores greater than or equal to the observed information score was used to calculate the probability of obtaining the observed information score by chance. A significance threshold of p < 0.01 was used to identify neurons with statistically significant spatial information (Deshmukh & Knierim, 2011).

#### Theta Phase Precession Analysis

Theta peaks in the local field potentials were detected as described by Deshmukh et al. (2010). Each spike was then assigned a phase (between 0° and 360°) using linear interpolation between consecutive peaks (Skaggs et al., 1996). For the circular track, 2D data was transformed into units of degrees on the track for linearized position estimates. Theta phase at which a place cell fired was plotted as a function of linearized position at which it fired to visualize theta phase precession as the animal passed through the place field.

#### Statistical Analysis

Two tailed tests were used for all quantitative statistical comparisons. Inter-frame intervals for both camera systems were normally distributed; Two-sample F-test for equal variances was used for comparing the two. Wilcoxon signed rank test was performed for all paired comparisons.

### Code Availability

Hardware setup instructions and video data acquisition codes are available at https://github.com/DeshmukhLab/PicameraPaper.

### Data Availability

The datasets generated during the current study are available from the corresponding author on reasonable request.

## Results

### Frames acquired using the Picamera system are temporally more stable compared to a commercial camera

Performance of the Picamera system was benchmarked against a commercial camera obtained as a part of the Neuralynx Cube-64 wireless recording system. Figure 2a shows the fraction of frames showing deviations from the expected inter-frame interval (IFI) for a video recorded for 4 hrs by the commercial camera and the Picamera in the same session. We defined jitter as the range of deviation from the expected IFI. The Picamera jitter of ± 0.025 ms was lower than the ± 7 ms jitter of the commercial camera. Thus, the Picamera system shows higher temporal accuracy over a long recording session compared to the commercial camera, giving two orders of magnitude improvement in jitter.

**Figure 2:**
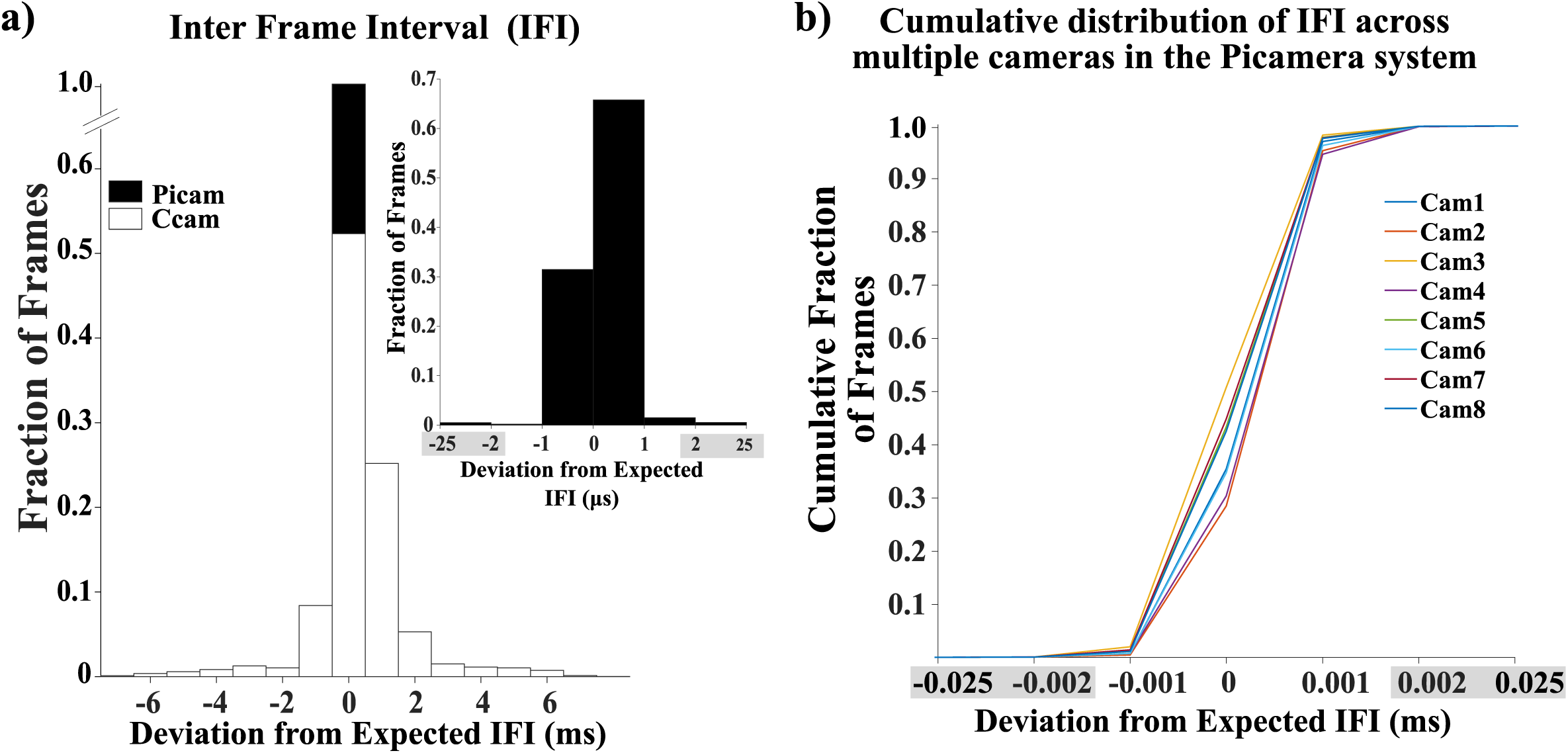
Frames acquired using the Picamera system show more regular inter-frame intervals (IFI) than a commercial camera. (a) Jitter in the IFI for a video recorded by the commercial camera (Ccam) and the Picamera system (Picam). In a 4 hr video recording session, the Picamera showed a jitter of less than ± 1 ms from the expected IFI vs ± 7 ms jitter shown by the commercial camera. The inset shows a zoomed-in plot for IFI for the Picamera showing a jitter of ± 0.025 ms. (b) All Picamera sub-units had a similar cumulative distribution with most of the frames lying within the ± 0.002 ms window for a 4 hr session. The plot shows that the IFI follows the same trend across cameras with each camera remaining extremely temporally accurate during a session. Since there is an extremely low fraction of frames outside ± 0.002 ms deviation, in place of the 0.001 ms bin width used for the inset histogram in (a) and cumulative distribution plot in (b), a larger bin width was used to combine data from ± 0.002 ms to ± 0.025 ms (marked by grey regions on the x axis to indicate substantially larger bin width compared to other bins on the same plot).

Similar IFI stability with ± 0.025 ms jitter was measured for videos recorded on eight Picamera sub-units simultaneously. The cumulative distribution plot for deviation from the expected IFI shows a steep increase at 0 ms as expected (Figure 2b). For all our recordings, all the frames across cameras were within ± 0.025 ms with > 98% of the frames lying within the ± 0.002 ms. The IFI distribution of the Picamera is statistically significantly smaller than that of the commercial camera (Two-sample F-test for equal variances, σ^2^_(commercial camera)_ = 3.1 ms, σ^2^_(Picamera)_ = 7.6 × 10^-7^ ms; f = 2.73 × 10^6^, df_(commercial camera)_ = 79909, df_(Picamera)_ = 95891, p < 0.0001). This temporal stability of the Picamera system can help accurately align frames across cameras with one another as well as the neural data and thus facilitate analysis of the neural and behavioral data at a higher temporal precision.

Since the data acquisition system uses a frame grabber to record video streaming from a camera, we tested whether webcams show a better temporal accuracy, as some behavior monitoring systems use webcams. We recorded videos using Logitech C170 USB webcam (Logitech, Lausanne, Switzerland) at 25 Hz. The webcam dropped an average of 8.26% of the frames giving us an extremely variable IFI (jitter = −28 ms to + 88 ms). Thus, webcams may not be ideal for use with neurophysiology systems.

### Reduced jitter improves estimate of neural correlates of behavior

We tested if the camera jitter affects our assessment of neural correlates of behavior by performing an explicit comparison of the Picamera and the commercial camera recording videos simultaneously with Cube-64 wireless transmitter recording neural activity. Multiple single units were recorded from the hippocampal formation while rats foraged for a food reward in different behavioral arenas (1 m × 1 m platform, 1 m diameter circular track, 1 m long linear track) across days. The animal’s position and head direction were estimated from the videos recorded using both the commercial camera and a single Picamera sub-unit. Spatial firing rate maps for units with significant spatial information score (spatial information > 0.25 bits/spike, p < 0.01 using rat position estimates from at least one of the two cameras) (n = 42) were generated for rat positions estimated from both the cameras used. Figure 3a shows firing rate maps for different cell types (two place cells, and one putative grid cell) recorded using the commercial camera and the Picamera. The place fields of spatially responsive neurons showed better tuning in the Picamera data than the commercial camera data. At the population level, place field size was significantly smaller for the Picamera data as opposed to the commercial camera data (Wilcoxon signed rank test, z = −3.88, p = 0.0001) presumably due to the greater temporal accuracy of the Picamera in positioning the animal (Figure 3b). These results motivated us to look for differences in spatial information and peak firing rate. As expected, peak firing rate (Wilcoxon signed rank test, z = 3.14, p = 0.0017), and spatial information (Wilcoxon signed rank test, z = 3.86, p = 0.00011) for the Picamera was significantly greater than the commercial camera, after Holm-Bonferroni correction for multiple comparisons (Holm, 1979).

**Figure 3:**
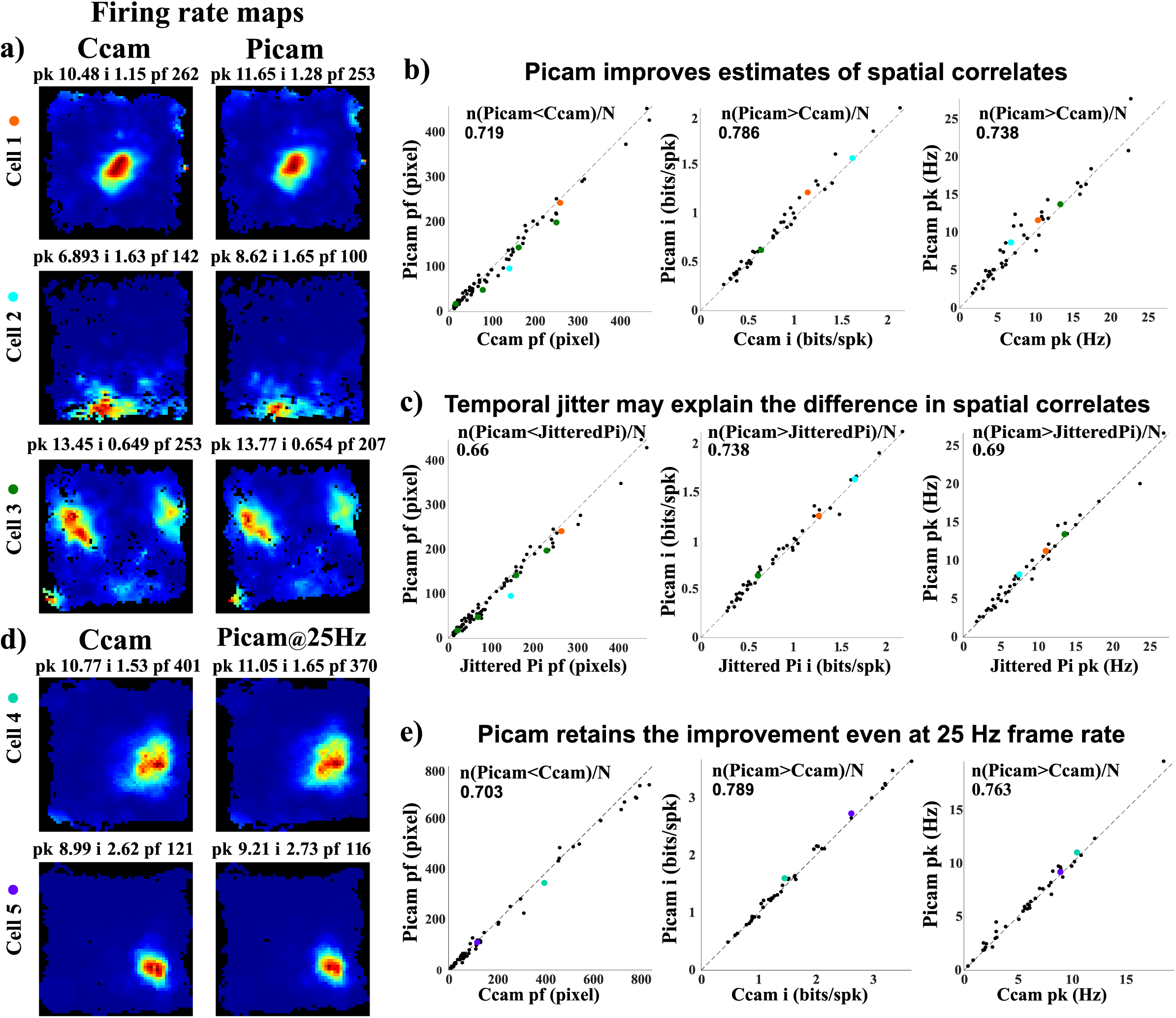
Estimates of neural correlates of space improve with enhanced temporal accuracy of video tracking. (a) Spatial firing rate maps of place cells (1^st^ and 2^nd^ row), and a putative grid cell (3^rd^ row) computed from videos recorded simultaneously on the commercial camera (left) and the Picamera (right). Peak firing rate (pk) in Hz, spatial information score (i) in bits/spike and place field size (pf) in pixels, for each rate map are presented at the top. Only the largest field size is presented for an individual cell (although other smaller fields are included in the analysis). For example, grid cell field size: commercial camera - 253 (other field sizes: 165, 80, 17) and Picamera - 207 (other field sizes: 149, 48, 12) (b) Scatter plots for place field size, spatial information, and peak firing rate for both cameras. Identity line (slope = 1) is marked on each plot. All place fields of a single cell meeting criteria in both the cameras were included in the analysis. Most of the points for place field sizes fall below the identity line indicating that the Picamera produces smaller place fields than the commercial camera on average. Correspondingly, most of the spatial information scores and peak firing rates fall above the identity line. (c) Scatter plots for place field size, spatial information and peak firing rate are calculated for the Picamera and Picamera data with frame times jittered using IFI distribution of the commercial camera (JitteredPi). Most of the points in the place field size scatter plot lie below the identity line indicating that the Picamera produces smaller place fields than the jittered Picamera on an average. Correspondingly, most of the spatial information scores and peak firing rates fall above the identity line indicating that the Picamera has higher spatial information scores and peak firing rate than the jittered Picamera on an average. (d,e) Picamera performs better than the commercial camera even at 25 Hz frame rate. (d) Spatial firing rate maps of place cells computed from videos recorded at 25 Hz simultaneously on the commercial camera (left) and the Picamera (right). (e) In agreement with (b) and (c), most of the points for place field sizes fall below the identity line indicating that the Picamera produces smaller place fields than the commercial camera on an average. Correspondingly, most of the spatial information scores and peak firing rates fall above the identity line indicating that the Picamera has higher spatial information scores and peak firing rate than the commercial camera on an average. The fraction of data points showing better performance with the Picamera is shown at the top of each of the scatter plots in (b), (c), and (e). Each cell shown in (a) and (d) was assigned a color which was then used to display that cell’s firing characteristics in the scatter plots in (b), (c), and (e).

We asked whether higher jitter in frame timestamp can lead to a large enough error in assignment of positions to individual spike timestamps to cause degradation of measures of spatial selectivity we compared above. We added jitter to the Picamera IFIs by sampling (with replacement) from the commercial camera IFI distribution, and generated rate maps using the jittered frame time estimate. There were significant reductions in spatial information (Wilcoxon signed rank test, z = 3.29, p = 0.001) and peak firing rate (Wilcoxon signed rank test, z = 2.99, p = 0.0028), and increase in place field size (Wilcoxon signed rank test, z = −3.13, p = 0.0018) from the Picamera to the jittered Picamera (Figure 3c). These results are consistent with the suggestion that higher temporal accuracy in the Picamera frame timestamps led to more accurate instantaneous position assignment and therefore improved measures of spatial selectivity.

We also tested whether the marginally higher frame rate of the Picamera (30 Hz) compared to the commercial camera (25 Hz) can explain the observed improvements by recording 38 single units at 25 Hz frame rate for both the systems in one rat. Consistent with the previous observations, the cells showed better spatial correlates with the Picamera. The Picamera showed smaller place field size (Wilcoxon signed rank test, z = −3.1, p = 0.002), higher spatial information score (Wilcoxon signed rank test, z = 2.87, p = 0.0041), and higher peak firing rate (Wilcoxon signed rank test, z = 2.53, p = 0.011) similar to the results shown earlier (Figure 3d,e). This continued better performance of Picamera despite reduction in frame rate indicates that the marginally higher frame rate of the Picamera (30 Hz vs 25 Hz) in the recordings above may not explain the improvements in the estimation of neural data.

Next, we tested if estimates of other neural correlates of behavior also improve with reduced jitter in the new system. For all 6 head direction modulated units in our dataset, the Picamera showed better estimates of head direction tuning (higher head direction peak firing rate and smaller full width at half maximum) (Figure 4a,b) compared to the commercial camera. Furthermore, the Picamera showed a tighter theta phase precession (O’Keefe & Recce, 1993) for both theta phase precessing place cells recorded on a circular track (Figure 4c).

**Figure 4:**
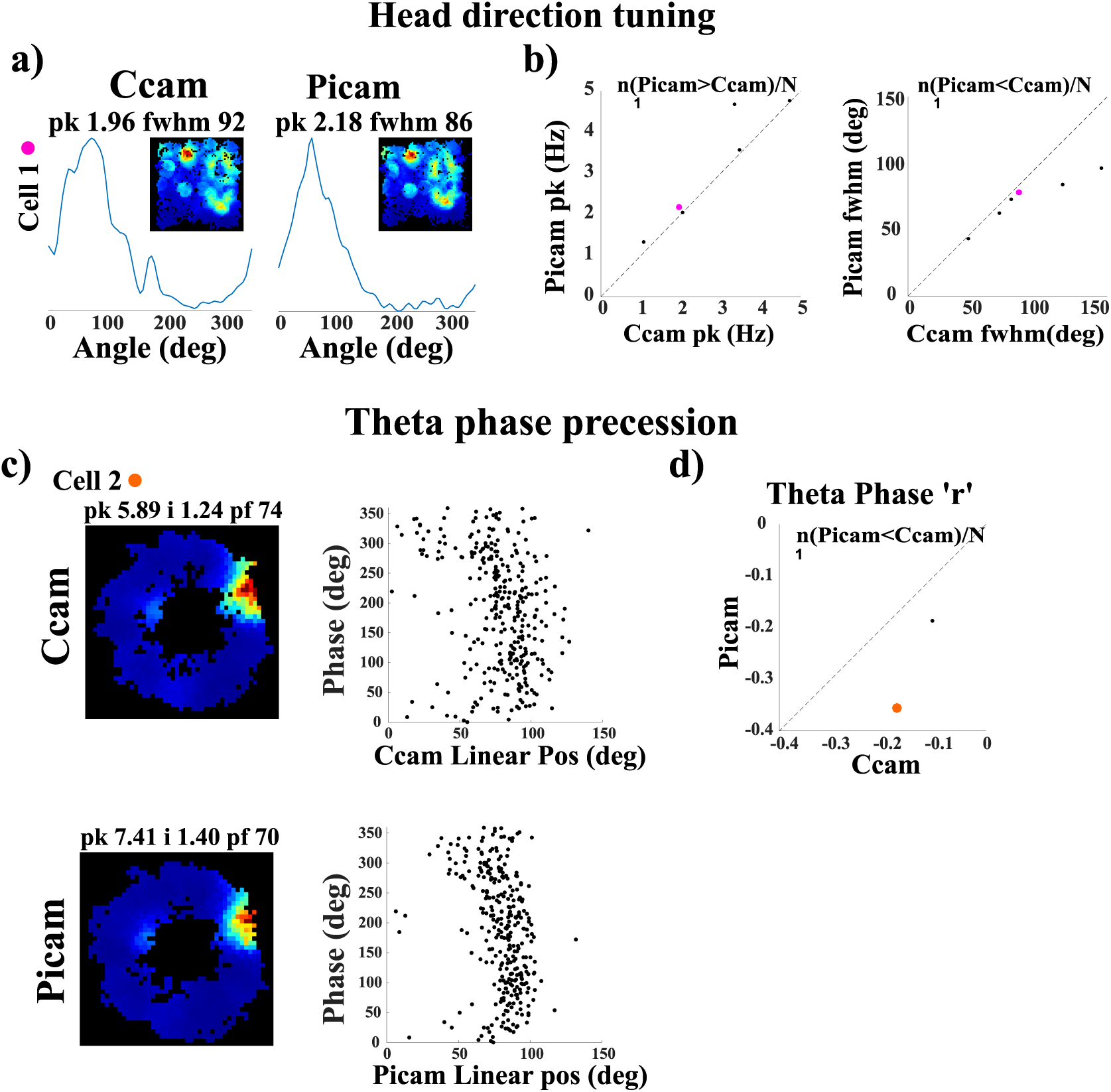
Estimates of head direction tuning and theta phase precession improve with enhanced temporal accuracy of video tracking. (a) Head direction tuning plot and spatial firing rate map (inset) for a head direction modulated cell computed from videos recorded simultaneously on the commercial camera (left) and the Picamera (right). Peak firing rate (pk) in Hz and full width at half maximum (fwhm) in degrees are mentioned at the top of the head direction plot. (b) The Picamera had higher peak firing rate and narrower full width at half maximum for all head direction cells (n = 6) compared to the commercial camera. (c) Spatial firing rate map and theta phase precession plot for a place cell computed from videos recorded simultaneously on the commercial camera (top) and the Picamera (bottom). (d) The Picamera had better theta phase precession as estimated by Pearson’s correlation coefficient (r) for both place cells recorded on a circular track compared to the commercial camera. The fraction of data points showing better performance with the Picamera is shown at the top of each of the scatter plots in (b) and (d). Each cell shown in (a) and (c) was assigned a color which was then used to display that cell’s firing characteristics in scatter plots in (b) and (d).

### Low jitter in individual Picamera frame timing allows precise synchronization across multiple cameras

The commercial camera has ∼ 40% of the frames outside 1 ms deviation from the expected IFI. Using multiple such cameras with their uncorrelated noise in frame timestamps can lead to higher chances of temporal misalignment of frames across cameras. This increased misalignment across cameras could worsen the estimates of neural correlates of behavior even more than that seen in the single camera case. The low jitter in IFIs of the Picamera sub-units discussed in previous sections predicted that their consecutive frames would be temporally closely aligned with frames from other Picamera sub-units, provided all the Picamera sub-units started recording videos nearly simultaneously. To test this prediction, we looked at how stable the inter camera frame interval stayed over the duration of the 4 hr recording session across 8 Picamera sub-units (Figure 5a). In multiple recording sessions, the starting times of the 8 cameras are within 0.5 ms of the first camera. Given the low IFI jitter across cameras, in multiple sessions, the consecutive frames from the 8 Picamera sub-units do not differ from that of the camera with shortest starting lag by more than 0.5 ms. The example in Figure 5a shows an across-camera frame time difference of less than 0.05 ms for all frames recorded over a 4 hr session. The extremely low inter camera frame timing difference shows that the entire system remained in sync during the recording session, facilitating temporal alignment of video frames across cameras at a sub-millisecond accuracy.

**Figure 5:**
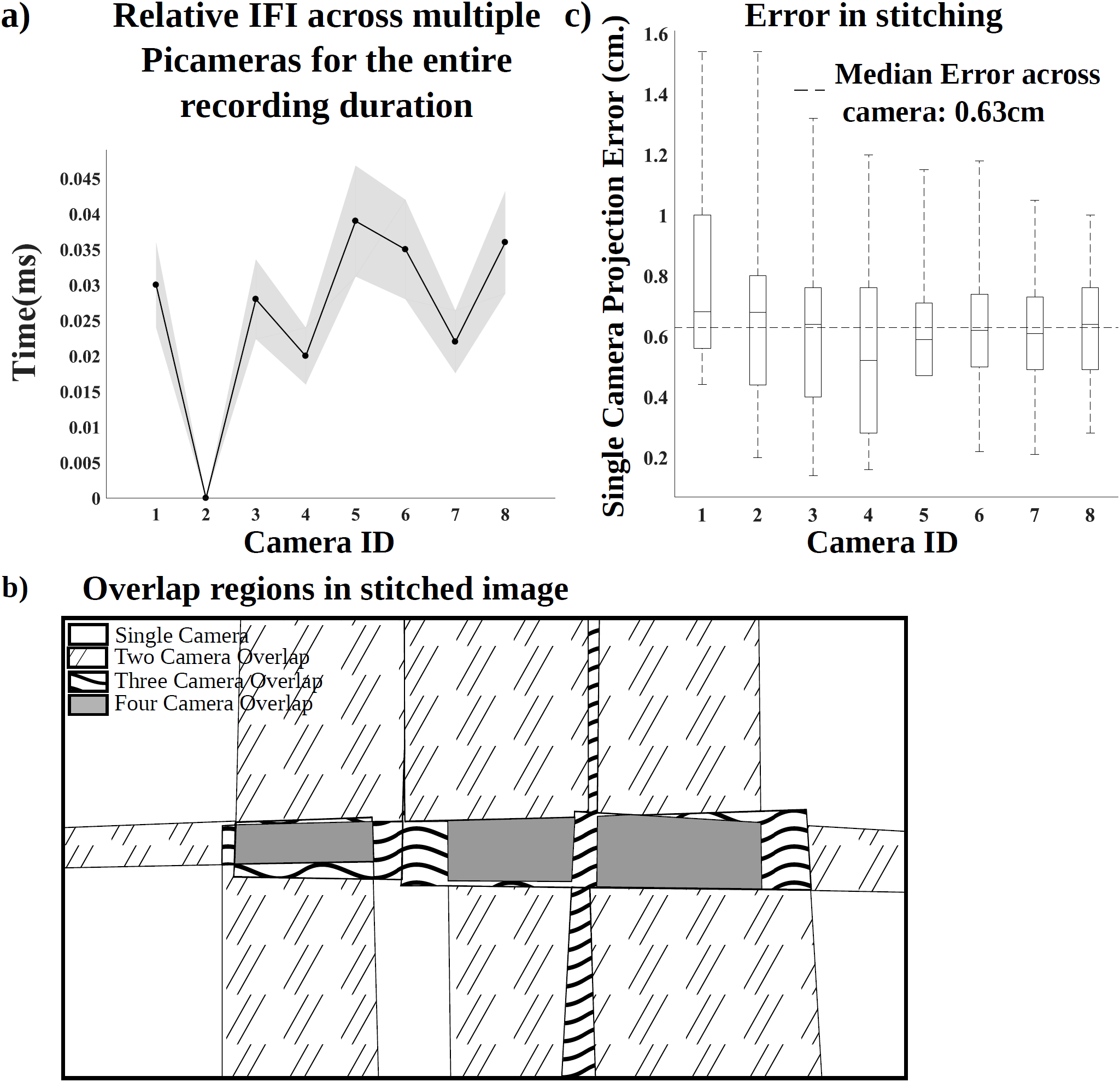
Synchronization across cameras and image stitching enable tracking of the animal in large environments. (a) The black line indicates the start frame time differences across cameras relative to the first camera to start recording the video (camera 2, here). Notice the spread of start times is within 0.05 ms after the first camera starts. The grey region indicates the relative frame time difference across cameras for all subsequent frames. (b) Top down view of the large room along with the regions covered by individual cameras after registering and aligning their images. Unfilled region, dashed line, wavy line and solid grey represents area of no overlap, overlap between 2 cameras, 3 cameras, and 4 cameras respectively (c) Projection error for each camera, defined as the distance between an intersection point formed after placing multiple strings to form a grid across the large room for that camera and its corresponding position across all cameras with fields of view overlapping at the intersection point. The central line, top edge, bottom edge, top bar and bottom bar of each box represents the 50^th^, 75^th^, 25^th^, 100^th^ and 0^th^ percentiles respectively of the samples. Dashed line represents median projection error across cameras. Maximum error noticed was 1.54 cm for the single camera and median error across cameras was 0.63 cm.

Using a video stitching algorithm, frames from individual Picamera sub-units were aligned in a single coordinate system which represented the entire maze. Figure 5b shows the overlapping regions across cameras after aligning them. The aligned frames were then checked for any distortions which could have been introduced because of the stitching algorithm, using an estimate independent of the calibration frames used for stitching. A grid was formed by stretching multiple strings across the length and breadth of the behavior arena which gave multiple intersection points between strings running orthogonally for each camera. When an intersection point is visible on multiple cameras, perfect alignment across cameras should place this intersection point at exactly the same x and y co-ordinates on the aligned frames of all the cameras. Thus, the difference in estimates from different cameras of positions of the shared intersection points provides a measure for accuracy of spatial location alignment using our stitching algorithm. We calculated the projection error for each camera, defined as the distance between an intersection point for that camera and its corresponding position across all cameras with overlapping fields of view. The maximum projection error for single camera with respect to others came out to be 1.54 cm and median error across cameras was 0.63 cm (Figure 5c). Thus, the stitched image has a spatial jitter of less than the pixel size (4 cm) used for creating firing rate maps of neurons when rats foraged in the large room.

### Tracking spatial selectivity of neurons from the hippocampal formation in a large room

Multiple Picamera sub-units were used to track position of a rat foraging in the large room. Videos recorded from each camera were aligned in a single coordinate system and the rat’s position was calculated after averaging position from transformed frames across cameras. Spatial firing rate maps were generated for neurons active during the behavior. Figure 6a shows trajectory plots and corresponding spatial firing rate maps for multiple place cells recorded in the 5.5 m × 3 m room and 2.75 m × 3 m room. Figure 6b shows head direction tuning curves for two head direction cells recorded in a 2.75 m × 3 m behavior room.

**Figure 6:**
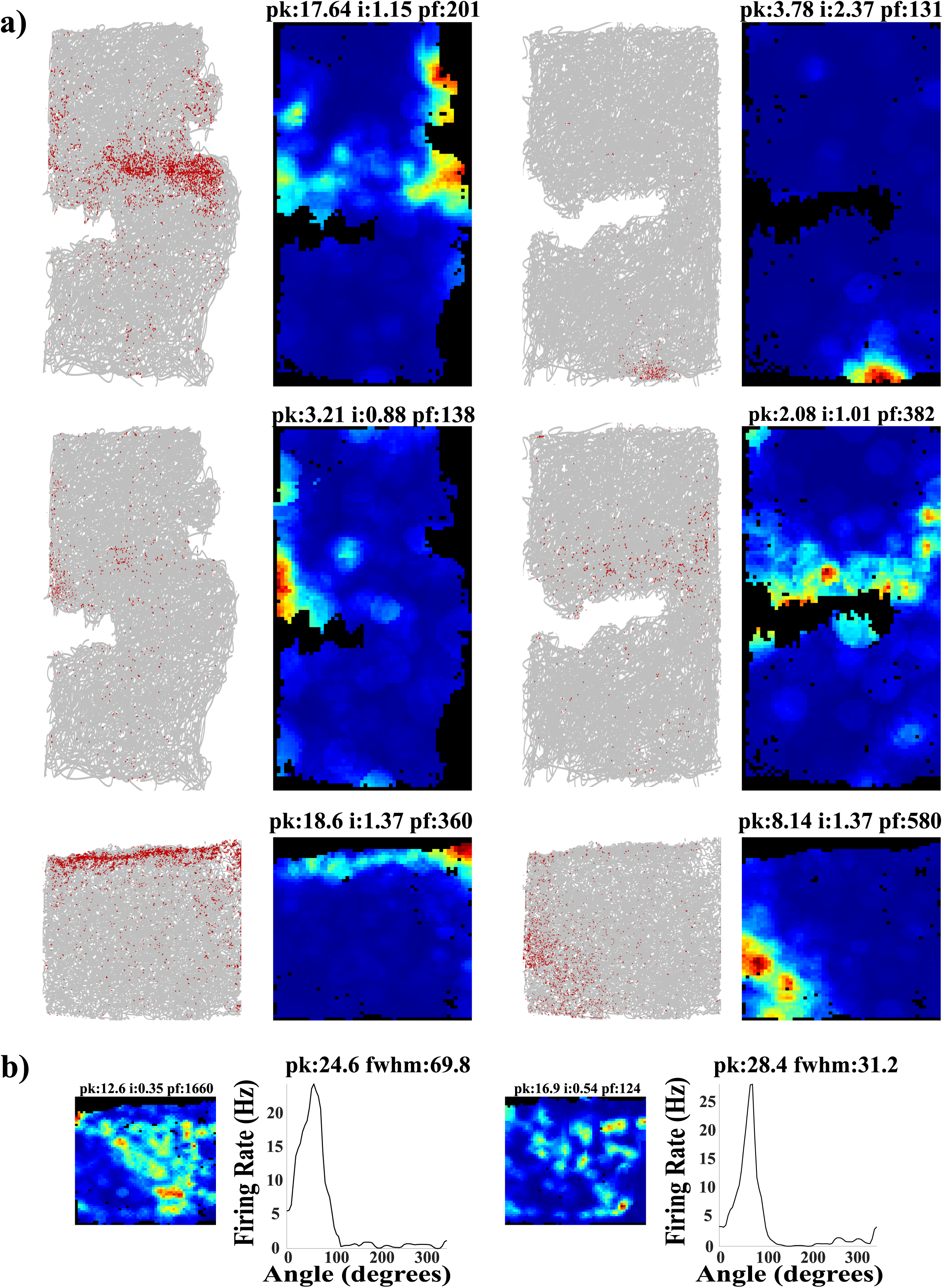
Neural representations of large spaces. (a) Trajectory plots and rate maps for place cells recorded in a 5.5 m × 3 m room and a 2.75 m × 3 m room. Grey lines in the trajectory plots mark the trajectory of the rat while red dots indicate locations where the neuron fired. (b) Rate map (left) and Head Direction Tuning curve (right) for two head direction cells recorded in a 2.75 m × 3 m behavior setup.

## Discussion

Extracellular recording studies from awake behaving rats are routinely used to understand the neural representations of space in the hippocampal formation. In the past, the sizes of behavioral arenas used in these studies have been typically constrained to up to 1 m^2^ by the weight/pull of the cable used to carry data from the animal to the amplifier/data acquisition system in the wired recording systems. In such small environments, a place cell typically has a single place field, enabling researchers to test how individual neurons respond to cue manipulations (Muller & Kubie, 1987; Shapiro et al., 1997; Fenton et al., 2000; Knierim, 2002; Huxter et al., 2003). Such responses could be interpreted as a dedicated code (Fenton et al., 2008), in which individual neurons encode unique spatial locations, and average of place field centers of neurons active at any given moment of time is sufficient to predict the actual location of the animal. In contrast, a number of studies have treated hippocampal code for space as an ensemble code, in which each location is represented by a population vector, defined as activity of a unique combination of neurons, associated with it (Wilson and McNaughton, 1993; Lee et al., 2004; Wills et al., 2005). The ensemble code is expected to perform better than the dedicated code in conditions under which a single place cell has multiple fields, as long as the population vectors at multiple locations where the given place cell is active are distinct from one another (Fenton et al., 2008). Explicit comparisons of hippocampal representations of small (68 cm and 76 cm diameter cylinders) and large (1.5 m × 1.4 m and 1.8 m × 1.4 m) environments reveals that the number of place fields/cell increase with increase in the recording area (Fenton et al., 2008; Park et al., 2011). Cells with multiple place fields create ambiguity in the dedicated place code but not in the ensemble code. Thus, the observation of multiple place fields in large environments argues in favor of ensemble place code over dedicated place code. In contrast, single place field seems to be the norm for place cells recorded on an 18 m long track (Kjelstrup et al., 2008). On an even longer, 48 m track, hippocampal neurons show a highly skewed (gamma-Poisson) distribution of propensity to be recruited to represent a spatial location, giving rise to a logarithmic distribution of number of place fields/cell, such that most cells have 0-2 fields but there are a few cells with 6 fields or more (Rich et al., 2014). This ensures availability of 5-20% place cells with single fields across arbitrarily large 1D environments, creating a possibility that a (quasi-) dedicated place code can be used in small as well as large environments. The discrepancy between the Fenton et al., (2008) and Park et al., (2011) studies on one side and Rich et al., (2014) study on the other side can be explained either as the difference between the mechanisms of encoding 1D and 2D environments, or as an effect of spatial scale. The second hypothesis predicts that as the spatial scale increases beyond the scales used in the 2D large space studies, the distribution of the number of place fields/cell will be skewed further, rather than shifted rightwards. Distinguishing between these two possibilities requires recording hippocampal activity from rats foraging in substantially larger spaces like the 16.5 m^2^ space we recorded from in this paper.

Recordings from larger spaces will also enable the experimenter to address a number of other questions relevant to our understanding of spatial representation at biologically realistic scales. For example, is the largest grid spacing limited to 1.7 m (Stensola et al., 2012, using 2.2 m × 2.2 m arena) or are there even lower resolution grid cells? How do environmental scale and geometry interact to distort grids (Stensola et al., 2015)?

The availability of wireless recording systems now facilitates recording of neural activity in large spaces. However, the commercially available extracellular electrophysiology systems still face limitations in terms of the number of cameras (usually one or two) used for tracking rats which constrains our ability to accurately track them in large environments. In this paper, we described a system for tracking rat behavior in large spaces. This Picamera system is temporally more accurate than a commercially available system used for benchmarking in this paper. It is easily scalable (due to its parallel architecture, adding more Raspberry Pi cameras to cover arbitrarily large areas is trivial) and can be adapted for use with complex environments with multiple occlusions. The Picamera system is cost effective because of the use of low-cost, off the shelf, easily available components (Prices from www.element14.com: Raspberry Pi 2 model B + Raspberry Pi camera module v1: $35 + $27 per unit, Arduino: $10) and makes the collection of high-quality, temporally accurate behavior datasets in large spaces feasible. Open-source libraries used in developing the code - OpenCV (used in stitching and position tracking) and Picamera Python library (used in acquiring videos and timestamps data) - made it possible to customize the code, and, in turn, allowed us to record videos at sub millisecond temporal accuracy.

Improvement in spatial and temporal accuracy of a tracking system is expected to lead to reduction in noise of estimation of behavioral variables like instantaneous position and head direction. This improved accuracy in tracking behavioral variables should lead to reduction in noise introduced by the tracking system in our estimates of spatial selectivity and head direction tuning of neurons. Predictably, the rate maps generated using the Picamera had significantly smaller place field size, increased peak firing rate and spatial information content as compared to a commercial system with higher temporal jitter. Similarly, the Picamera showed sharper head direction tuning as well as tighter theta phase precession compared to the commercial system. Understanding of mechanism and functions of theta phase precession will benefit from increased accuracy in quantifying theta phase precession. (J. O’Keefe & Recce, 1993) showed that hippocampal place cells fire in earlier and earlier phases of theta oscillations in the hippocampal local field potentials as the rat traverses through their place fields. This relationship between theta phase and position improves the estimate of the animal’s position over firing rate alone (Jensen & Lisman, 2000; Reifenstein et al., 2012). While linear (O’Keefe & Recce, 1993) or circular-linear (Huxter et al., 2008; Kempter et al., 2012) regression is routinely used to quantify theta phase precession, the observed dynamics of theta phase precession are more complex (Skaggs et al., 1996; Yamaguchi et al., 2002; Mehta et al., 2002; Souza & Tort, 2017), with the rate of precession changing as a function of location within the place field. Different proposed mechanisms for theta phase precession predict different shape of theta phase precession and variability of the preferred theta phase at different locations within the place field (reviewed in (Burgess & O’Keefe, 2011). Improved accuracy of the position tracker will clearly enhance our estimate of the shape as well as variability of theta phase precession, enabling us to distinguish between the different models. Trial by trial analysis of theta phase precession (Schmidt et al., 2009; Reifenstein et al., 2012; Feng et al., 2015; Souza & Tort, 2017) will benefit even more, given that a single aberrant spike would have a larger effect on the single trial estimate. Similarly, estimates of theta phase precession in 2D environments (Skaggs et al., 1996; Huxter et al., 2008; Jeewajee et al., 2014) will show improvement with better position tracking. The Picamera system with its sub-millisecond temporal accuracy is well suited for such applications, even at current frame rate of 30 Hz, but this can further improve at higher frame rates. While the current system is capable of acquiring at 60 Hz with minimal (0.31%) frame drops and sub-millisecond accuracy, our preliminary tests indicate that the newer version of the hardware (Raspberry Pi 3 with Raspberry Pi camera module v2) is capable of acquiring at 100 Hz at sub-millisecond temporal accuracy without dropping frames.

Correlated with the debate about the exact shape of theta phase precession, there is an ongoing debate about the exact shape of the place field. The shape of the place field is critical to the two competing models of theta phase precession: while the ramp excitation model requires asymmetric place field with a slow increase in firing rate as the rat approaches the location with peak firing rate and a rapid fall off in firing rate as the rat exits the place field (Mehta et al., 2002), the oscillatory interference model requires a symmetric place field ((J. O’Keefe & Recce, 1993); (Burgess & O’Keefe, 2011). While the criteria used for defining the place field can affect the estimated asymmetry, precise position tracking, and the consequent increased confidence in the exact shape of the place field can further help resolve this question.

We demonstrated our ability to record from large spaces by characterizing neural activity in 5.5 m × 3 m room using the Picamera system with 8 sub-units (Figure 6). The stitching and position estimation algorithm are completely automated and only require the registration data to be calculated once beforehand. Our stitching approach gives a maximum error of 1.54 cm in estimating position of the rat. This error is less than the resolution used for generating spatial firing rate maps. Thus, the Picamera system along with the wireless recording system can now be used to perform experiments in larger sized as well as complex environments with occlusions (e.g. burrows).

In summary, the system described in the present work satisfies all the criteria desirable in an efficient tracking system for use with large and complex environments: easy to use, adaptable, temporally accurate, low-cost, scalable, open-source and easily available. The system overcomes the bottleneck on tracking animal behavior in large spaces, therefore reducing the gap between the natural environments and experimental setup.

## Acknowledgements

We thank Nishita VS, Indraja R. Jakhalekar, Manish Mohapatra and Mekhala Kumar for support in data collection; Deepa Jain and Lou Blanpain for help with early prototypes of a multicamera tracking system; Aditi Bishnoi for help with collecting 25 Hz Picamera data; Sushant Kuchankar for assistance with TTL wiring setup; James J Knierim, Francesco Savelli, and Vyash Puliyadi for comments on the manuscript. This work was supported by Wellcome Trust/DBT India Alliance Grant IA/S/13/2/501024

---

**Supplementary Figure 1:**
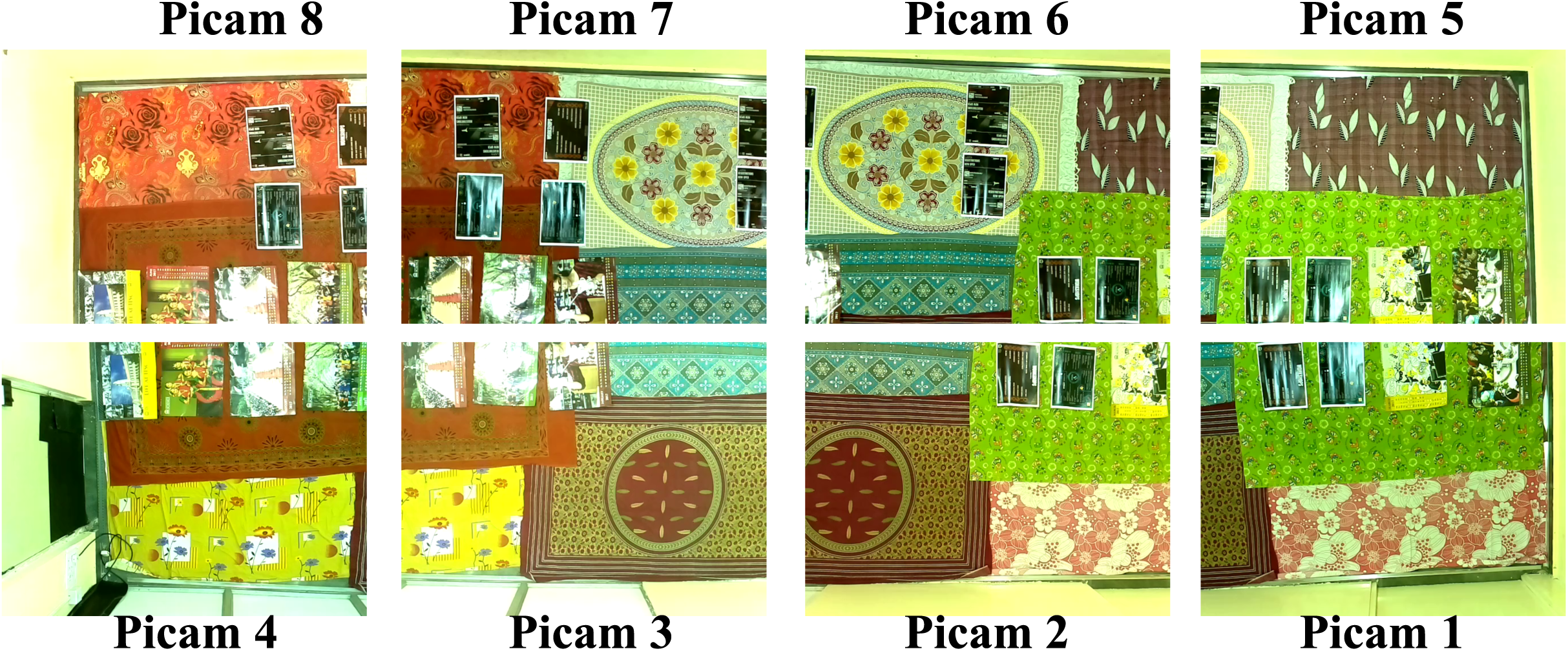
Calibration frame from each camera: Bedsheets with multiple unique identifiable features were placed in the large room and an image (calibration frame) was acquired on each Picamera sub-unit. These features were then used for calculating registration data for each camera.

